# Long-term near-continuous recording with neuropixels probes in healthy and epileptic rats

**DOI:** 10.1101/2023.02.16.528689

**Authors:** Antoine Ghestem, Marco N Pompili, Matthias Dipper-Wawra, Pascale P Quilichini, Christophe Bernard, Maëva Ferraris

**Author notes:** Correspondence should be addressed to MF or MNP. Equally contributing first author (Antoine Ghestem, Marco N Pompili, Matthias Dipper-Wawra). Equally contributing last author (Pascale P Quilichini, Christophe Bernard, Maëva Ferraris).

## Abstract

Neuropixels probes have become a crucial tool for high-density electrophysiological recordings. Although most research involving these electrodes is in acute preparations, some scientific inquiries require long-term recordings in freely moving animals. Recent reports have presented prosthesis designs for chronic recordings, but some of them do not allow for probe recovery, which is desirable given their cost. Others appear to be fragile, as these articles describe numerous broken implants. This fragility presents a challenge for recordings in rats, particularly in epilepsy models where strong mechanical stress impinges upon the prosthesis. To overcome these limitations, we present a new prosthesis specifically designed to protect the probes from strong shocks and enable the safe retrieval of electrodes after experiments. This implant was successfully used to record from healthy and epileptic rats for up to three weeks almost continuously, with a probe retrieval and reuse success rate of 91%, improving previously described recycling performances.

## INTRODUCTION

Recording the activity of many neurons over extended periods is essential for understanding physiological and pathological processes that can evolve over minutes to decades. For instance, learning complex tasks and memory consolidation can take from a few days to several weeks (Frankland and Bontempi, 2005). Brain networks are continually remodeled by both external factors, such as life experiences, and internal ones, like circadian rhythmicity. Indeed, the molecular architecture of brain networks, its activities and its functions change during the night and day cycle (Brüning et al., 2019; Noya et al., 2019; Debski et al., 2020). Moreover, appreciating the dynamic evolution of brain networks on a timescale even slower than the circadian one is particularly relevant in the context of neurological disorders that can develop over months to years, such as epilepsy and Alzheimer’s disease. In epilepsy, seizure probability displays a periodicity that can vary from several days to several months (Karoly et al., 2021). Thus, understanding physiological and pathological processes requires following brain activity over long periods in freely behaving animals.

Neuropixels 1.0 probes, which have 960 recording sites along a 10 mm shank, and version 2.0 with four shanks and a total of 5120 recording sites, allow multiple brain regions to be recorded simultaneously along the probe axis (Jun et al., 2017, Steinmetz et al., 2021). Many laboratories have used these probes in acute, head-fixed experiments with various behavioral tasks (e.g. Allen et al., 2019; Bennett et al., 2019; Evans et al., 2018; Jun et al., 2017; Kostadinov et al., 2019; Musall et al., 2019; Vélez-Fort et al., 2018). However, there are fewer examples of chronic recording in unrestrained animals (Krupic et al., 2018; Gardner et al., 2019; Böhm and Lee, 2020). Yet to understand the neural mechanisms underlying cognition it is essential to track behavior and neuronal activity in longitudinal studies in freely moving animals (Pompili and Todorova, 2022), ideally in ethologically relevant settings (Demars et al., 2022).

Although several studies proposed approaches for chronic recordings with Neuropixels probes in unrestrained rodents (Juavinett et al., 2019; Luo et al., 2020; Steinmetz et al., 2021; van Daal et al., 2021), we encountered two issues that remained to be addressed. First, some rat implants were not designed to recover the probes (Steinmetz et al., 2021), despite their high cost (approximately €1000), while it would be advantageous to have the ability to reuse them in order to decrease the cost of experiments. Second, proposed implant designs appeared fragile with many broken implants, particularly when the recordings were performed in rats (Juavinett et al., 2019; Luo et al., 2020; van Daal et al., 2021). This issue is particularly relevant when performing recordings in epileptic animals, whose generalized motor convulsions and jumping expose the implants to particularly strong mechanical stress and shocks.

To address both issues, we designed a new implant that allows retrieving and reusing the probes and is specifically designed to resist strong shocks to record epileptic rats. Here, we first demonstrate how to build and surgically implant this prosthesis for chronic recordings. Then, we describe the procedures that we used to record in a near-continuous 24/7 manner (12 h recording blocks separated by a few minutes) in freely moving control and epileptic rats in their home cages. With this protocol, we successfully recorded tens of neurons for weeks and explanted the probes in 91% of the animals, allowing us to reuse them. We observed only one breakage, substantially improving previously described recycling performances.

## MATERIALS AND METHODS

All references of the chemicals, materials and tools (software and hardware) are detailed at the end of this paper (**Supp. Tab. 1**).

**Table 1.** Weight of the 3D printed parts of the prosthesis.

| 3D-printed parts | Weight (g) |
| --- | --- |
| Left-side enclosure | 3.68 |
| Left-side enclosure with hex nuts, copper tape and ground wire | 4.58 |
| Right-side enclosure | 3.12 |
| Right-side enclosure with copper tape | 3.83 |
| Headstage lock | 0.40 |
| Headstage lock with hex nuts | 0.45 |
| Probe guide | 0.06 |
| Probe guide with hex nuts | 0.11 |
| Foothold | 0.13 |
| Lid | 1.45 |
| Full weight of the design with all screws and the electrode/headstage | 13 |

### 3D designing and printing

We designed all parts with Tinkercad (Autodesk, CA, USA), which were 3D printed with a Form 2 printer (Formlabs) with a resolution of 50 µm and Grey Pro Resin (Formlabs, MA, USA). After printing, we cured them with UV light at 80°C for 15 minutes (Form cure, Formlabs). All design files are available on our GitHub repository (https://github.com/INS-PhysioNet/npx-ghestem).

### Design overview and assembly instruction

The part of the implant not mechanically connected with the probe is made of four printed pieces: two enclosures, a headstage lock (HL) and a lid (**Fig. 1A-C**). The enclosures are fixed to the skull to protect the probe (see surgery procedure) and their interior is lined with copper tape to provide shielding from electrical noise. Their rostral parts are flat in order not to obstruct the visual field of the animal. A hole in the left-side enclosure allows the electrical shield to be connected to the recording ground: a screw (00-90 1/2”) is fixed with a brass hex nut (00-90) through the hole and soldered to a wire with a pin connector (tulip pin ic socket) (**Fig. 1A, D**).

**Figure 1:**
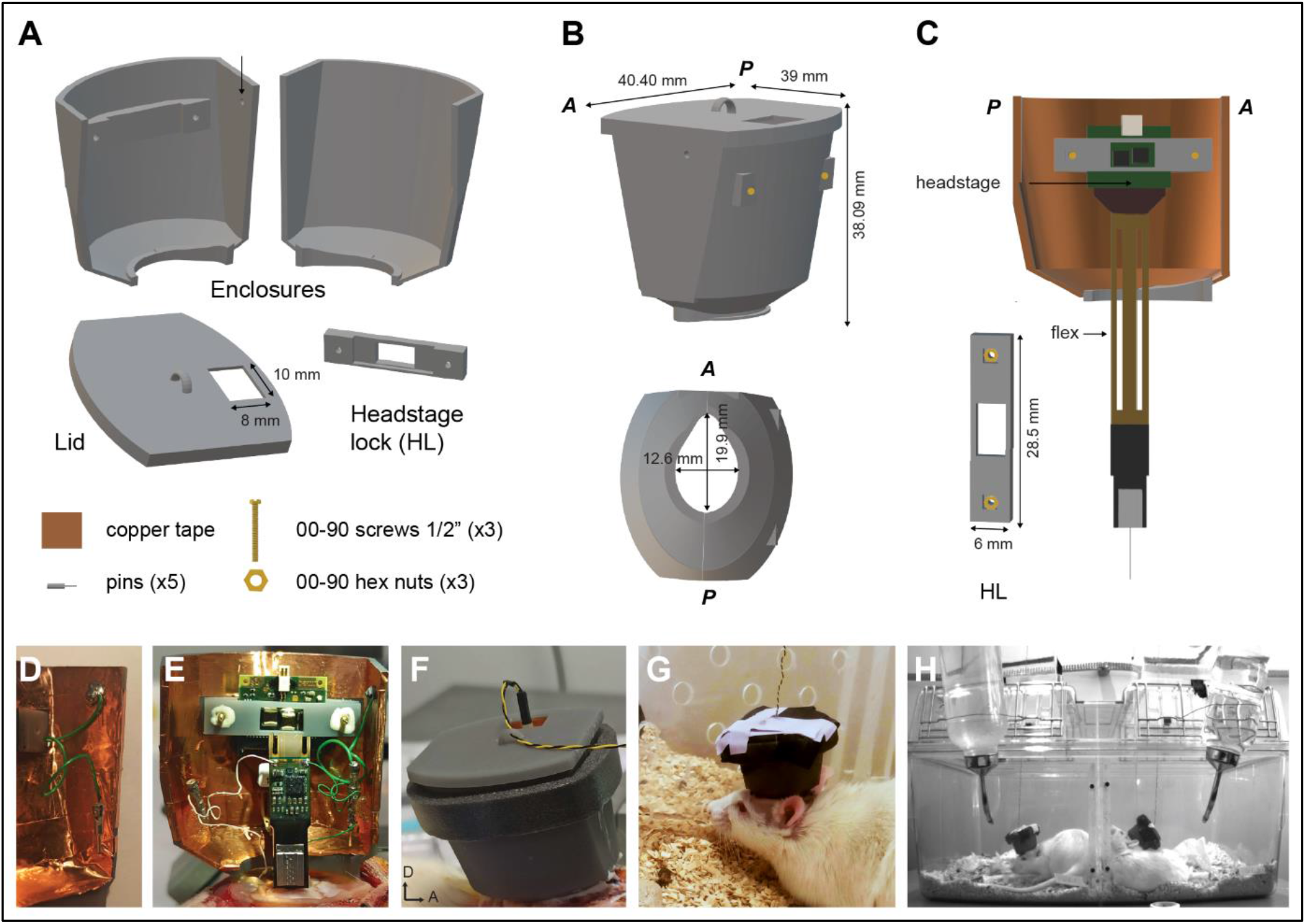
Implant design overview. ***(A)*** The part of the implant not mechanically connected with the probe is composed of four 3D printed pieces: two enclosures, a lid, and a headstage lock (HL). In addition, three hex nuts, three screws, five pins, and copper tape are required to assemble the implant. Note the designed hole (arrow) in the left-side enclosure. ***(B)*** Dimensions of the implant. ***(C)*** Schematic of the headstage connected to the probe and attached to the left-side enclosure with the HL. ***(D)*** Photograph of the ground wire attached to the left-side enclosure prepared before surgery. ***(E)*** Photograph of a fixed Neuropixels probe and its headstage during surgery. Note that both ground (green) and reference (white) wires are connected with pins to their relative screws above the cerebellum. The ground wire is also connected to the Faraday cage. ***(F)*** The interface cable can be connected to the headstage without removing the lid. The lid hook (visible at the centre of the lid) limits the tensions on the cable. An adhesive foam tape surrounds the implant to cushion mechanical shocks. ***(G)*** Rat chronically implanted in its home cage. The lid is secured with tape. Note the transparent separation with holes splitting the cage in two compartments for social interactions. ***(H)*** Rats simultaneously recorded in their large, double compartment cage. A: anterior, D: dorsal, P: posterior.

The prosthesis is large enough to accommodate the headstage, thus preventing potential mechanical damages caused by repeated connections. During the surgery procedure, the headstage is connected to the probe and secured on the left-side enclosure (**Fig. 1C,E**). For this purpose, the HL is equipped with two holes accommodating two brass hex nuts (00-90) fixed with dental cement (Super Bond) (**Fig. 1C**). This method allows the headstage to be effortlessly secured in the enclosure with two matching brass screws (00-90 1/2”).

A small rectangular window on the lid allows the connection of the animal to the acquisition system without opening the cap. A small hook at the center of the lid helps secure the cable with a piece of tape, limiting the tension applied to the headstage connector due to the animal’s movements and preventing damages. After connecting the animal, the window is taped shut and the animal can be recorded in its home cage (**Fig. 1F-H**).

In order to secure the probe to the skull during the surgery, the probe is attached beforehand to a holder composed of two 3D printed parts: the probe guide and the foothold (**Fig. 2A**). First, two brass hex nuts (00-90) are fixed to the probe guide with dental cement (Super Bond). Then, using a binocular, the probe guide is cemented (Super Bond) to the back of the probe while the probe remains attached to its holder (provided with the probe by Imec, Löwen, Belgium) (**Fig. 2B**). Finally, the probe guide is slid and secured to the foothold with two screws (00-90 3/32”; **Fig. 2B-C**). To allow connection to the reference and ground channels, two small wires with tulip pin connectors (ic socket) are soldered on the relative contacts on the probe pads.

**Figure 2:**
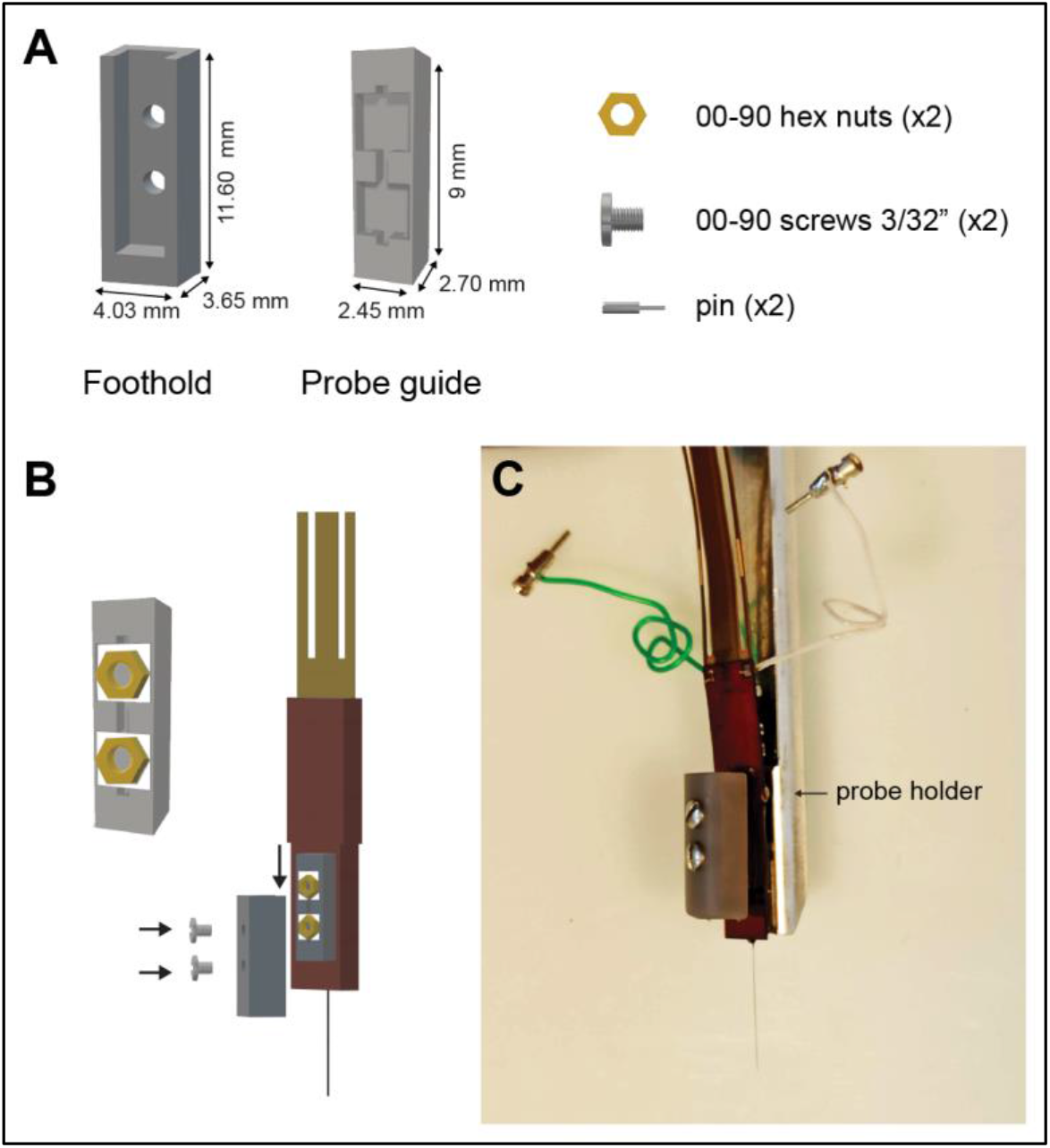
Neuropixels probe pedestal system. ***(A)*** The probe is mounted on a stand made of two 3D printed pieces: a probe guide and a foothold, needing two hex nuts, screws, and pins. ***(B)*** The foothold is screwed to the probe guide, which itself is fixed at the back of the probe. ***(C)*** Photograph of a mounted probe. Note the ground (green) and reference (white) wires of the probe.

The pedestal holding the probe allows its easy removal at the end of the experiment since only the foothold is fixed to the skull. The experimenter only needs to unscrew the probe guide from the foothold to retrieve it (see procedure for recovering the probe). To facilitate the removal of the probe at the end of the experimental procedure, vaseline is applied on the foothold before the assembly is made.

Once assembled, the prosthesis has a total weight of 13 g (**Tab. 1**)

### Subjects

All experiments were conducted in accordance with the guidelines of Aix-Marseille Université, Inserm, and local ethics committee. The French Ministry of University and Research (approval numbers 01451-02 and #20325-2019041914138115 v2) approved the protocol use for this study. The rats (adult male Wistar Han, n = 11) were kept on a 12-hour light/dark cycle with free access to food and water. Animals were housed in pairs in large cages (61.2 cm × 43.5 cm × 21.6 cm) divided in two compartments by a transparent partition (Plexiglas) with holes (diameter: 11 mm) allowing social interactions (**Fig. 1G-H**), and were handled daily and familiarized to the experimenter for at least 10 days before the surgery. Status epilepticus (SE) was induced in nine rats by a single intraperitoneal (i.p.) injection of pilocarpine (320 mg/kg) at 7 weeks (∼250 g) (Levesque et al., 2016). To reduce peripheral effects, rats were pre-treated with methyl-scopolamine (2 mg/kg, i.p.) 30 min before the pilocarpine injection. SE was stopped by diazepam after 60 min (2×10 mg/kg, i.p., two doses within a 15-min interval). Then, rats were hydrated with saline (2×2 ml, subcutaneous injection, 2 h and 4 h after the first diazepam injection). On the following days, they were fed with a porridge made of soaked pellets until they resumed normal feeding behavior. Probes were implanted >5 weeks following SE induction.

### Surgery procedure

To induce anesthesia, rats were placed for a few minutes in an induction chamber with sevoflurane (8%, 1 l/min O2), until the breathing rate slowed down. They were then placed on a heating pad (Harvard Apparatus, MA, USA) and anesthesia was maintained with sevoflurane through a ventilation nosepiece (4%, 0.5 l/min O2). Heart rate, breathing rate, pulse distension, and arterial oxygen saturation were monitored with a pulse oximeter (MouseOx, STARR Life Sciences, PA, USA) during the whole surgery to ensure stable anesthesia and monitor vital constants. Analgesia was provided with a subcutaneously (s.c.) injection of buprenorphine (0.025 mg/kg). We evaluated the correct anesthesia level by the absence of paw retraction reflex. Then, the head was secured in a stereotaxic frame (Kopf, CA, USA) and an injection of lidocaine (5 mg/kg, s.c.) was made along the midline of the scalp for local anesthesia after shaving the fur. A midline incision was made to expose the skull, which was cleaned before marking the position of the craniotomy for the probe with a pencil. Next, the lateral temporal muscles were carefully detached from the skull with a scalpel and four holes were drilled (two per side) where four miniature stainless-steel screws (00-90 3/32”) were inserted to serve as anchors for the implant (**Fig. 3A**). Two additional screws (00-90 3/32”) soldered to pin connectors (tulip pin ic socket) were placed above the cerebellum to serve as ground and reference electrodes (**Fig. 3A**). Any bleeding from the skull, skin or muscles was cauterized. A ring of dental cement (Paladur, Kulzer GmbH, Germany) was applied covering the ground/reference and anchor screws and extended until the anterior limit of the incision (**Fig. 3B**), serving as the base for anchoring the implant. Particular care was given to ensure that its thickness was even across its entire perimeter and that ground and reference wires were oriented inward.

**Figure 3:**
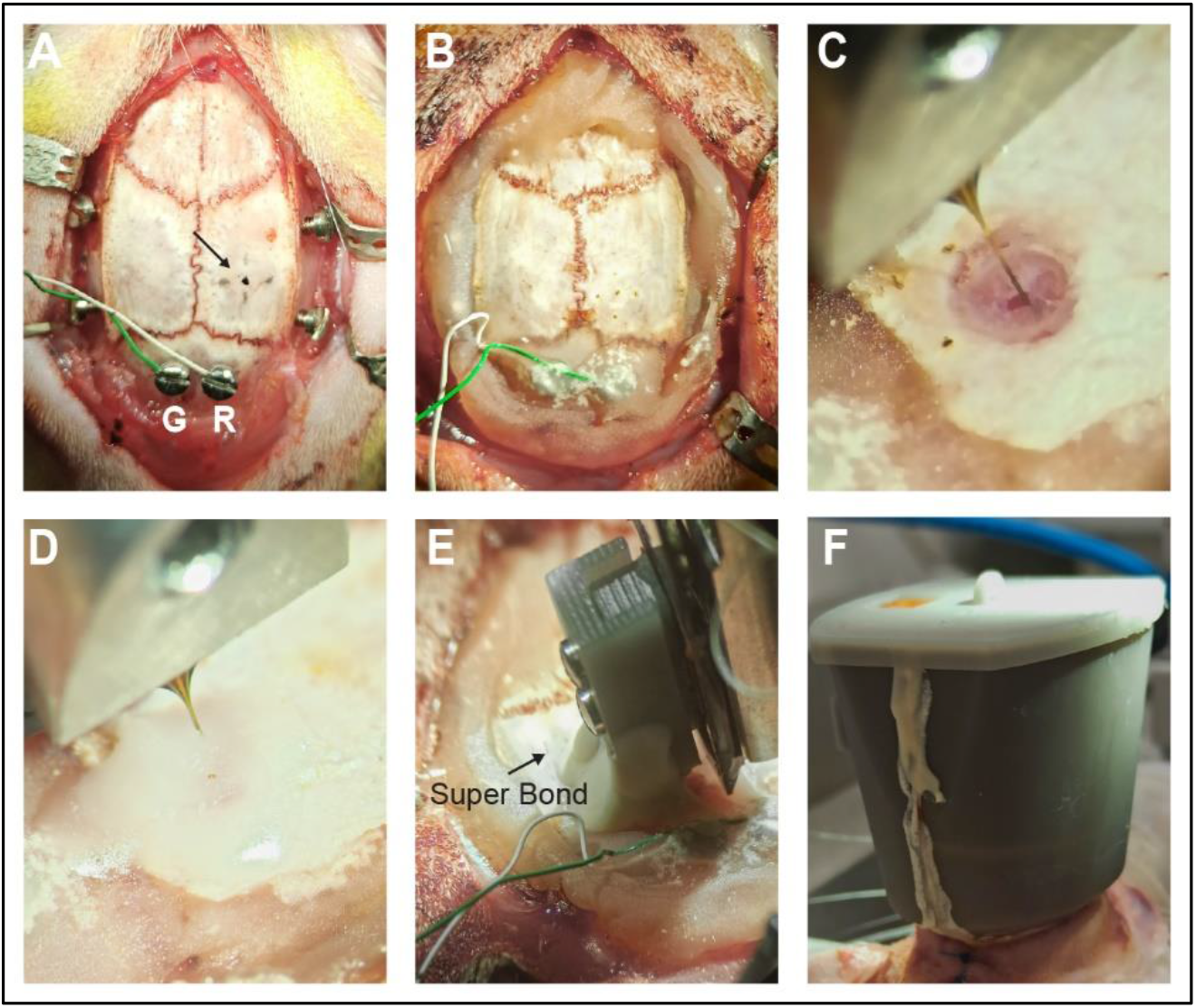
Probe implantation. **(A)** The two screws on each side of the skull and above the cerebellum serve as anchors, ground (G), and reference (R) electrodes, respectively. Note the marked position (arrow) of the craniotomy. ***(B)*** A ring of dental cement covers the screws and builds the platform on which the implant is mounted. ***(C)*** The probe is inserted into the center of the craniotomy after removing the dura-mater. ***(D)*** The craniotomy is sealed with silicon and a paraffine/wax mix and ***(E)*** dental cement is applied on the foothold to fix the probe. ***(F)*** Fully mounted implant at the end of the surgery. Note the presence of dental cement along the junction between the two enclosures. The skin is closely sutured with one or two stitches around the implant.

Next, a craniotomy was performed, and the dura-mater was carefully opened with a hook-shaped needle (**Fig. 3C**). Saline solution was applied to prevent drying of the brain surface. The Neuropixels probe was placed on its holder and DiIC18(3) (Interchim, France) was carefully applied on the back of the probe (i.e. the side without recording sites) so that its position could be determined with *post hoc* histological analysis.

The holder was mounted on a motorized manipulator (IVM single, Scientifica, United Kingdom) on the stereotaxic frame and the ground and reference wires were connected to their corresponding wires on the probe with the pins. Then, the probe was lowered in the brain at a constant speed of 2.5 µm/s to reach the target structures. For our recordings we aimed to record both dorsal and ventral parts of the hippocampal formation and areas along the track, either targeting the hippocampus (n= 7; AP: -4.68; ML: -3.1; DV: -9.6; angle: 5.5°) or the subiculum (n= 4; AP: -5.88; ML: -2.3; DV: -9.5; angle: 11°) through the thalamus to the amygdala. The headstage was connected to the recording system and electrophysiological activity was continuously monitored to ensure proper positioning of the probe relative to electrophysiological landmarks. To implant the probe as much as possible and still being able to seal properly the craniotomy, probe insertion was temporarily stopped 1.5 mm before reaching the final coordinates.

An anti-inflammatory drug (dexamethasone 0.1%) was applied in the craniotomy, which was then sealed to protect the brain with low-viscosity silicone and covered with a mixture of paraffine and wax melted with a cauterizer (**Fig. 3D**).

Once the probe was fully inserted to its target location, a dental resin adhesive cement (Super Bond C&B, Sun Medical, Japan) was prepared and gently applied with a needle or a brush between the bottom of the foothold and the skull (**Fig. 3E**). Once the cement had set, the probe holder was removed. The headstage was then secured on the left enclosure of the cap with the flex cable softly curled in an S-shape (**Fig. 1C,E**). This enclosure was then positioned over half of the dental cement ring with the stereotaxic arm and cemented to it (Super Bond, C&B). Once the cement set, ground pins were connected to the cap for electrical shielding and the second enclosure was cemented using the same procedure.

To tight the seal between the two sides of the implant’s enclosures, the gap between its two sides was also sealed with dental cement (Super Bond C&B; **Fig. 3F**). Finally, ground and reference pins were glued inside the cap with epoxy at two different locations to ensure their insulation. If necessary, the most rostral and caudal sections of the scalp incision were sutured to ensure a tight seal between the prosthesis and the skin. The lid was then placed, and adhesive foam tape was placed directly underneath to absorb eventual shocks (**Fig. 1F**). The lid was secured with tape and a small piece of tape was used to close the window for the connection (**Fig. 1G**). Finally, the rat was removed from the stereotaxic frame and received subcutaneous injections of saline solution (2 ml twice to avoid dehydration), analgesic (buprenorphine, 0.05 mg/kg), and antibiotics (sulfamethazine and trimethoprim, 10 mg/kg) treatments. On the following two days, the rat received subcutaneous injections of antibiotics (sulfamethazine and trimethoprim, 10 mg/kg, s.c) and analgesic (meloxicam, 1 mg/kg).

### Probe Recovery procedure

At the end of the recordings, we anesthetized the rats with a mixture of ketamine/xylazine (ketamine: 100 mg/kg, xylazine: 10 mg/kg). First, we opened the implant along the contact line between the two enclosures using cutting pliers, then the enclosure not holding the headstage was removed after cutting its basis with cutting pliers. Then, the headstage was unscrewed and the enclosure on which it was mounted was detached as with the other. The rat was then placed in a stereotaxic frame to secure its head and the Neuropixels holder was mounted on a motorized manipulator and carefully aligned with the Neuropixels probe, considering the angle between the main axis of the probe and the animal’s skull. The probe was attached on the holder and the screws on the back of the foothold were removed and then was slowly retrieved using the motorized manipulator (**Fig. 4A-B**). The probe was afterwards rinsed with distilled water, placed in a 1% Tergazyme solution for one hour for cleaning, gently brushing the probe with small bits of wet wiper (Kimwipes, Kimtech), and finally rinsed with distilled water the following day.

**Figure 4:**
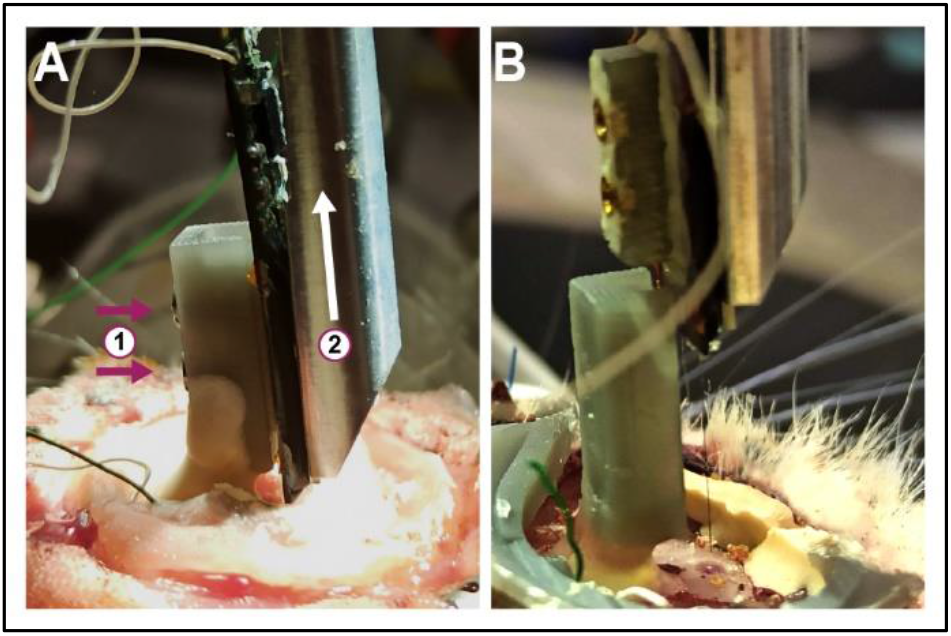
Probe recovery. ***(A)*** To retrieve the probe at the end of experiment, (1) the probe is first unscrewed from the foothold and (2) slowly lifted with the micromanipulator. ***(B)*** Overview after the probe was removed from the brain while the foothold remains fixed to the skull.

### In vivo electrophysiological recording and data storage

Extracellular signals were recorded simultaneously from 2 animals (**Fig. 1H**) with a commercially available Neuropixels acquisition system combined with National Instruments PXI hardware (Putzeys et al., 2019). A pulley system and counterweight lifted the interface cable out of the animal’s reach, following its movements. The recording system was controlled with Open Ephys software (Neuropixels PXI plugin) using the external reference connected during surgery. The software wrote data on disk on two separate binary files: one containing the wide-band signal (0.3-10 kHz bandwidth, 30 kHz sampling rate, 250x gain), and the other lower band signal (0.5-500 Hz bandwidth, 2.5 kHz sampling rate, 125x gain) for analysis of the local field potential (LFP).

Recordings were performed in ∼12 h blocks generating for each animal ∼1.8 TB of raw data per day. In parallel with recordings, data blocks were moved to a redundant dual-server storage system. Each storage server is connected via PCI-E to SAS interface cards (Dell HB 330) to multiple bays (Dell MD1200) of 12 spinning SATA hard disks. Each disk exists in two physically separate redundant copies on the two servers (located in distinct rooms), allowing any individual drive or entire server to fail or be destroyed without loss of data.

### Spike sorting

The 12 h blocks of data recorded on a given day were concatenated using the dedicated Neurosuite (Hazan et al., 2006) function. We used Kilosort v3.0 (Pachitariu et al., 2023) for data preprocessing and spike sorting with default parameters. To reject noise and to merge erroneously split units, we visually inspected and manually curated the resulting clusters with Phy (https://github.com/cortex-lab/phy). We rejected units as noise either if their waveform clearly did not correspond to an action potential, or if the waveform was identical on all the channels. Units were rejected as excessively noisy if they presented excessive refractory period violations in their autocorrelograms. We inspected in phy all the pairs of units labelled as *good* by Kilosort and merged those that showed a similarity index > 0.9. We also those corresponding to different spike types of complex spike bursts (Ranck, 1973; Harris et al., 2000) of the same unit. Finally, we retained only the units that were stable across the whole 24 h sessions (i.e. which did not suddenly appear or disappear).

### Seizure detection and classification

In pilocarpine-treated animals, epileptic seizures were systematically detected in a semiautomated manner using a custom-made Matlab algorithm based on the band power of the LFP recording (whole band). Each detected electrophysiological event was visually verified. The stage of the epileptic seizure was then determined in the video recording based on Racine scale (**Supp. Vid. 1**; Racine, 1972).

### Histological Analysis

After electrode retrieval, animals were injected with a lethal dose of pentobarbital (150 mg/kg, i.p.) and perfused intracardially with a 4% paraformaldehyde solution in 0.12 M phosphate buffer (PB), pH 7.4. After removal, we fixed the brains for 1 h at room temperature in the same PFA solution and then rinsed them in PB before cryoprotecting them in a 20% sucrose PB solution overnight and freezing them on dry ice. The brains were sliced into 40 µm-thick coronal sections using a cryostat. The position of the electrodes was revealed by the presence of DiIC18(3), which had been applied on the back of the electrodes before insertion and confirmed histologically on fluorescent Nissl staining.

Fluorescent Nissl staining was realized (Neuro-Trace 500/5225 Green Fluorescent Nissl Stain, Invitrogen, MA, USA) on free-floating sections. Sections were soaked in a KPBS 0.02 M solution containing 0.1% triton X-100 during 10 min then rinsed twice in KPBS 0.02 M. They were subsequently transferred into the Neuro-Trace (1/200 in KPSB 0.02 M) for 20 min in the dark. Finally, slices were soaked again in KPBS 0.02 M + 0.1% triton X-100 and washed three times in KPBS 0.02 M. Section were mounted on glass slides and coverslip with Fluoromount-G (Electron Microscopy Sciences, PA, USA).

## RESULTS

### Recording in blocks of 12 h allows untangling of the interface cable

To test our protocol, we implanted 11 rats to record from both the dorsal and ventral parts of the hippocampal formation and nearby areas, targeting either the dorsal and ventral hippocampus or the dorsal and ventral subiculum, therefore also recording the thalamus and the amygdala (**Fig. 5A-B; Tab. 2**). The animals were recorded in pairs using double-compartment cages with a transparent wall with holes separating the two compartments to allow the two rats to maintain social contact (**Fig. 1G-H**). This is particularly important for epileptic rats, who tend to develop aggressive behavior and display a high level of stress when singly housed without social contact (Manouze et al., 2019). Such animals become very difficult to handle, which becomes an issue when it is necessary to connect and disconnect the interface cable. During continuous long-lasting recordings (12h in the present experiments), the cable tends to tangle, especially at night when rats are more active. As a result, the cable extension becomes shorter, which limits animals’ movement and increases the risk of disconnection and/or damage. Our solution was to untangle the cable twice a day by briefly interrupting the recording and disconnecting the rat from the recording system, which resulted in ∼12 h recording blocks.

**Figure 5:**
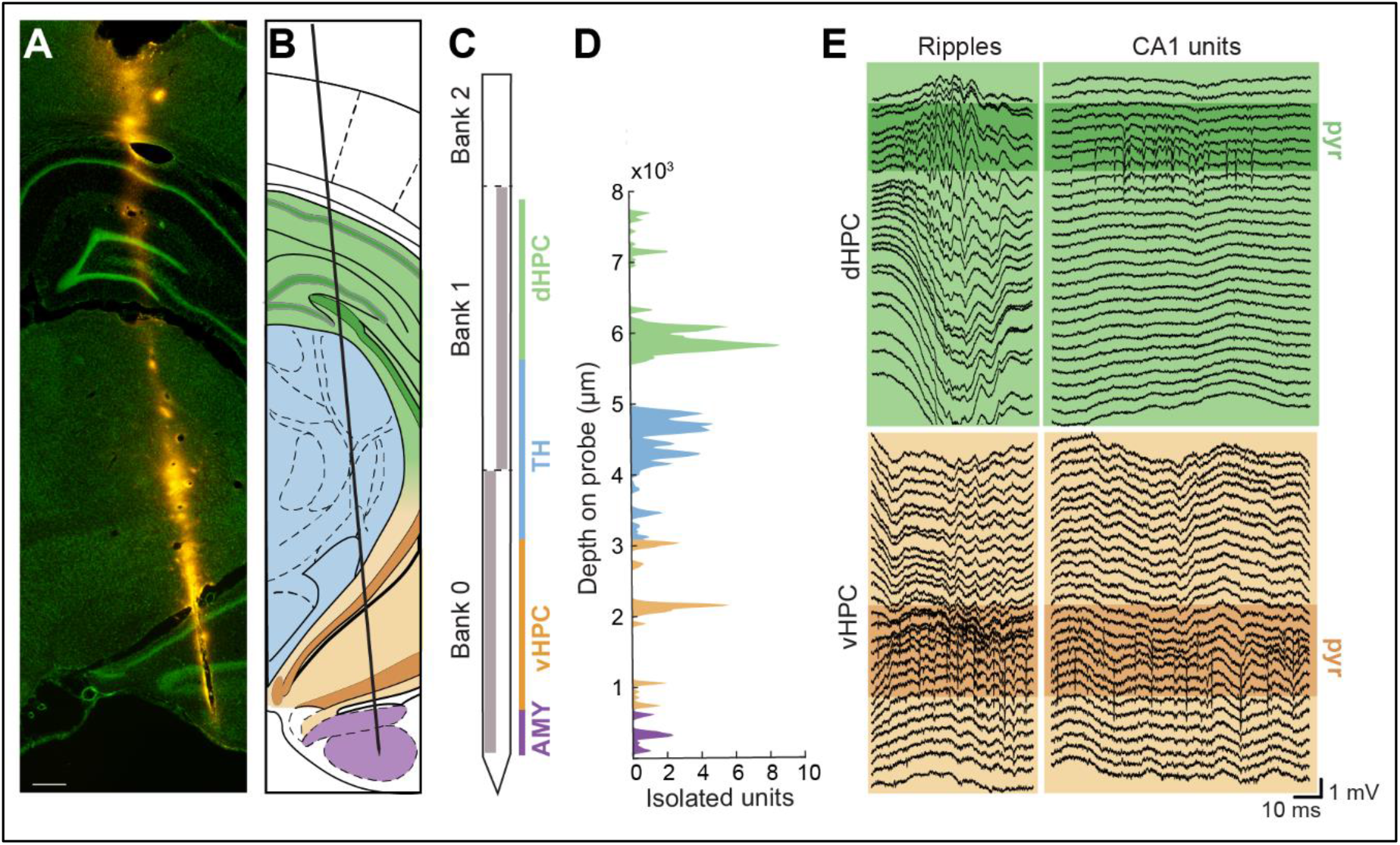
Recording across the whole dorso-ventral axis of the hippocampal formation in freely behaving rats. ***(A)*** Location of the Neuropixels probe marked with DiIC18(3) (orange) over a green, fluorescent Nissl staining. Scale bar: 400 µm ***(B)*** Neuropixels probe implantation trajectory (coordinates from bregma AP: - 4.68; ML: -3.1; DV: -9.6; angle: 5.5°) through the dorso-ventral axis of the hippocampus according to the Paxinos-Watson atlas. ***(C)*** Schematic of the probe and of the selected channel banks (grey). ***(D)*** Number of isolated units recorded in rat #2 relative to the probe depth (total, n = 350). ***(E)*** Representative example of simultaneous dHPC and vHPC CA1 pyramidale layer (pyr) recordings during both slow wave sleep and wake. AMY: amygdala, dHPC: dorsal hippocampus, TH: thalamus, vHPC: ventral hippocampus.

**Table 2.** Overview of Neuropixels probe implantation experiments. Rats were implanted either aiming the hippocampus (HPC; AP: -4.68; ML: -3.1; DV: -9.6; angle: 5.5°) or the subiculum (SUB; AP: -5.88; ML: -2.3; DV: - 9.5; angle: 11°) along the dorso-ventral axis. Epileptic rats are marked with an asterisk.

| Rats | Structure | Implanted period | Probe | Outcome | Probe serial # |
| --- | --- | --- | --- | --- | --- |
| #1 | HPC | 15 days | New | Intact | 19301415172 |
| #2 | HPC | 16 days | New | Intact | 19301415482 |
| #3* | HPC | 24 days | New | Intact | 19301403532 |
| #4* | HPC | 11 days | New | Intact | 19301414031 |
| #5* | HPC | 17 days | Recovered | Intact | 19301415482 |
| #6* | SUB | 16 days | New | Intact | 19301415562 |
| #7* | HPC | 15 days | New | Intact | 19301415632 |
| #8* | SUB | 14 days | Recovered | Intact | 19301415172 |
| #9* | HPC | 16 days | Recovered | Intact | 19301415632 |
| #10* | SUB | 14 days | Recovered | Intact | 19301415172 |
| #11* | SUB | 36 days | Recovered | Broken (lost) | 19301415172 |

### Stable yield of well-isolated units across days in healthy and epileptic rats

We recorded from 384 channels by selecting half of the channels from bank 0 and 1 to cover the entire hippocampal formation traversed by the probe from dorsal to ventral parts (**Fig. 5C**). Over the course of 1-3 weeks, we recorded the activity from different regions, including the hippocampus/subiculum, thalamus, and amygdala (**Fig. 5C-E**). We then concatenated the 12 h recording blocks into ∼24 h sessions and used Kilsort3 for spike sorting. After manual curation, we isolated ∼9-69 units per session in one control and two epileptic rats.

In epileptic rats, we detected a lower number of well-isolated units, nevertheless these figures remained relatively stable within each rat and session, despite the occurrence of 7 and 8 epileptic seizures (stage 3-5), respectively (**Fig. 6A, Supp. Vid. 1**). We did not observed degradation of the signal after seizures events (**Fig. 6B**).

**Figure 6:**
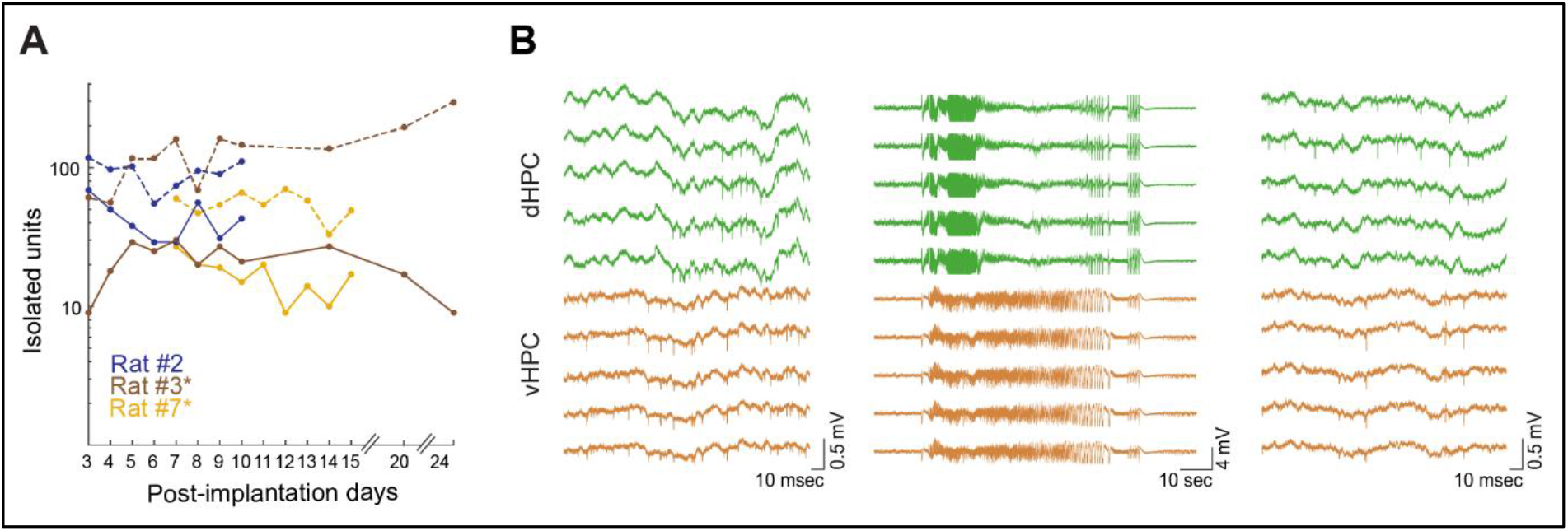
Recording stability in healthy and epileptic rats. ***(A)*** Number of isolated units before (dashed line) and after manual curation (solid line) across days in control (#2) and epileptic rats (#3*, #7*). Both epileptic animals had seizures during recordings. ***(B)*** Representative examples of wide band signals in both dorsal and ventral hippocampus (rat #3*) ∼20 minutes before (left) and ∼5 minutes after (right) a bilateral tonic-clonic seizure (middle). Note the presence of units also shortly after the seizure.

It is important to note that manual curation after spike sorting appeared necessary to reject noise. In both healthy and epileptic rats, the number of units automatically classified as “good” by Kilosort was higher (median: 3.2, min/max: 1.7/33.8) than after manual curation which rejected on average 66 % of the units (**Fig. 6A**).

### Successful retrieval and reuse of the probes relies upon a perfect sealing of the craniotomy during surgery

Rats were implanted for up to 36 days (17±2), after which probes were explanted (**Tab. 2**). Only one epileptic rat, with severe and numerous epileptic seizures lost his prosthesis 36 days after implantation (rat #11). Rat #11 had 51 seizures during 14 recorded days while rats #3* and #7* whose data is shown in Figure 6A had 7 and 8 seizures each in 22 and 9 days, respectively. In all other animals, no probe was found broken inside the enclosure meaning that the latter was strong enough to protect the probe (**Tab. 2**). Moreover, in all these animals we were able to explant functional Neuropixels probe (**Fig. 7A**), therefore showing that our design and surgical protocols are particularly suitable for functional probe recovery for reuse. To avoid seeping of liquid, we sealed the craniotomy before finishing the insertion of the probe in the brain to prevent cerebrospinal fluid leakage. The only critical step during the probe removal was to carefully align the holder with the probe (**Fig. 4**).

**Figure 7:**
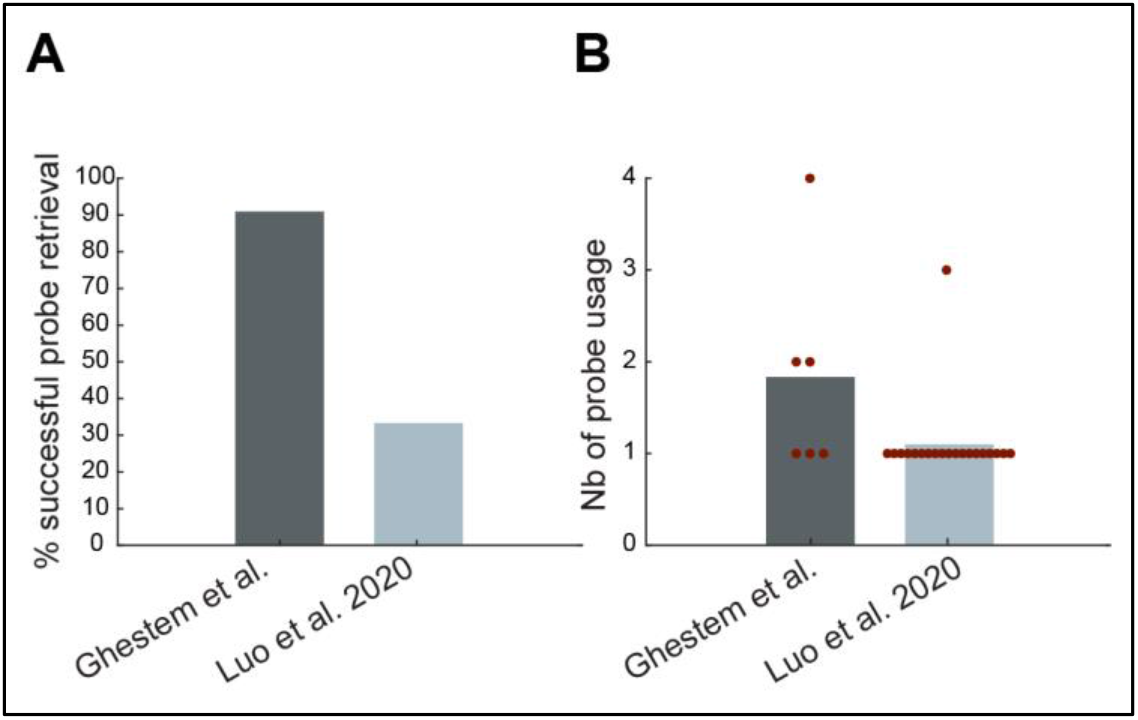
Probe explantation and reuse performance compared to previously described rat implant design. Percentage of probes successfully retrieved ***(A)*** and reused ***(B)*** in comparison to a previously described rat chronic implant allowing probe recycling (Luo et al. 2020).

The explanted probes were successfully reused in 6 rats, with one probe used for 4 different implantations and two for 2 (**Fig. 7B**), thus greatly enhancing previously described recycling performances (Luo et al., 2020). Overall, we used six probes to record 11 animals.

## DISCUSSION

Here, we introduce a novel protocol that employs a 3D printed prosthesis for chronic implantation and reuse of Neuropixels probes in freely moving rats, both epileptic and control. Our implant is specifically designed to withstand the strong mechanical stress and shocks encountered during recordings in epileptic animals, making it more solid than previously presented designs. Additionally, our system allows for the easy explantation of the probes at the conclusion of the experiment. By utilizing our design, we have achieved a higher success rate in probe recycling compared to previous reports, resulting in reduced overall research costs.

Our experiments show that the implant design effectively protects the probe from any impact across multiple weeks, even for epileptic animals. We successfully recorded nine epileptic animals with epilepsy over up to 3 weeks in a near-continuous manner. Although some animals had several seizures, the design of the implant prevented probe damage during repeated head shocks in all but one animal (rat #11). Indeed, in contrast to other systems (Juavinett et al., 2019; Luo et al., 2020; van Daal et al., 2021), the external part of the implant is not directly attached to the Neuropixels probe or its pedestal, preventing external impacts from being transmitted to the probe therefore moving and potentially breaking it.

Importantly, the protocol described here greatly increased previously described performances in recycling of the probes. The key factor for successful probe recover is the sealing of the craniotomy during the surgical implantation. Indeed, significant infection in the craniotomy results in the seeping of the cerebrospinal fluid that dries in the foothold, sticking the probe to the tissue, hence preventing its extraction without breakage.

In our experiments, we recorded the activity of an average of 26 ± 2.7 neurons per session for a period of 1-3 weeks. Overall, these numbers are lower than those reported in other studies (Juavinett et al., 2019; Luo et al., 2020; van Daal et al., 2021), especially for the two epileptic rats. Multiple factors could account for these results. First, previous studies primarily recorded from the cortex and midbrain (Jun et al., 2017; Juavinett et al., 2019; Luo et al., 2020; van Daal et al., 2021), where cell bodies are larger and more evenly distributed, resulting in a higher probability of isolating single units. In contrast, the thalamic nuclei have a lower density of smaller cells. In the hippocampus, the pyramidal layer is densely packed with cells but only ∼80-100 microns thick, which means that only 6-10 channels of a Neuropixels probe cover the layer. Additionally, the pilocarpine model of epilepsy is associated with significant cell loss in different subfields and layers of the hippocampus, as well as with a global structural reorganization of its networks (Dinocourt et al., 2003; Boulland et al., 2007). Second, in previous reports, the density of recording electrodes was maximized by using the electrodes from only one bank of the probe (Juavinett et al., 2019; Luo et al., 2020; van Daal et al., 2021), which increased the number of units recorded. In our study, we aimed to record along the whole septo-hippocampal axis of the hippocampal formation; therefore, we selected sparser channels from two banks, halving the density of recording channels. Finally, in our study, we worked with 24-hour-long sessions, whereas previous studies sorted single units from recordings lasting 5 minutes to 3 hours (Jun et al., 2017; Juavinett et al., 2019; Luo et al., 2020; van Daal et al., 2021). Luo et al. (2020), in particular, primarily studied the stability of spiking signals in 10-minute recordings without manual curation. They also found that manual curation reduced the number of isolated units. We replicated this observation and primarily studied only manually curated data, making direct comparison with other studies using uncurated results impossible. It is also worth noting that none of these studies used Kilosort3 as a spike sorting software, but rather Kilosort, Kilosort2, or JRCLUST (Jun et al., 2017; Juavinett et al., 2019; Luo et al., 2020; van Daal et al., 2021), and in some cases, they also used different probe versions (Jun et al., 2017; Juavinett et al., 2019).

Here, we chose to take advantage of the length of the probe to record simultaneously dorsal and ventral regions of the hippocampal formation and perform recording as stable as possible across days. To this aim, we did not mount the Neuropixels probe on a movable microdrive but rather fixed it on the skull. Nevertheless, other target regions and experimental aims might require adapting our design to use a microdrive. This can be done by substituting the 3D printed probe guide cemented to the back of the probe with a microdrive (such as the Cambridge Neurotech nanodrive) and assembling it with a similar foothold system.

Finally, our protocol allowed us to perform recording on an almost continuous basis in freely moving rats. Such a protocol opens new possibilities to investigate the dynamics of neuronal activity over long time spans. In experimental epilepsy, in particular, it is important to record continuously to assess seizures and interictal activity (Karoly et al., 2021). In this context, our protocol offers a new scale of recording compared to telemetry systems typically used in experimental epilepsy but only equipped with a few channels (Manouze et al., 2019; Debski et al., 2020). Although the recording of our rats took place in their home cages, our system also allows recordings during behavioral tasks. The connection of the rat can be done without the removal of the lid and thus with minimum handling and stress for the animal. An improvement to our system would be the use of a rotative cable commutator, to avoid cable entanglement and thus brief disconnection of the rat between recording blocks. Nevertheless, the only currently available system, produced by SpikeGadgets (CA, USA) requires the use of their specific headstage and implant, which do not allow the separation between the probe pedestal and the enclosure, necessary for epileptic animals. Moreover, full continuous recordings will also require the development of novel algorithms for offline spike sorting of months long recordings from Neuropixels probes.

In conclusion, our design provides a new tool for continuous recording of freely moving rats, pending the development of wireless system with high channel count whose battery can be recharged online. Our design allows the recycling of Neuropixels probes across multiple animals and thus significantly reduces the cost of experiments. We share all of 3D designs to help standardize chronic procedures and allow other laboratories to replicate them.

## ACKNOWLEDGMENTS

This project was funded with a grant from the Agence National de la Recherche (grant # ANR-19-CE14-0036 STRESS) to CB. A Walter Benjamin Fellowship from the Deutsche Forschungsgemeinschaft (DFG, #455053935) supported MDW.

## AUTHOR CONTRIBUTIONS

Research project conception and design: CB, MF, AG, PPQ. Funding gathering: CB. Technological implementation: AG, MNP, MDW. Experiments: MF, MDW, AG. Data processing and analysis: MF, MNP, MDW. Visualization and writing: MF, MNP with comments from all authors. Project supervision and management: MF, CB, PPQ, MNP.

## SUPPLEMENTARY INFORMATION

**Supplementary Video 1. Spontaneous seizure**. Representative video of a stage 5 seizure of rat #3 (on the left side) displaying chewing and facial movements, subtle head nodding, forelimb clonus, rearing and falling. See Fig. 6B (middle traces) for the corresponding electrophysiological signals.

**Supplementary Table 1.** Materials list.

| Resources | Designation | Company | Reference |
| --- | --- | --- | --- |
| Drug/Chemical compound | Buprenorphine, Buprecare 0,3mg/ml injectable solution. | Axience | GTIN 03760087151244 |
|  | Dexamethasone, Maxidex 0.1% eye drops | Novartis | Cip : 3400930652473 |
|  | Dental cement, Paladur liquid | Kulzer | 64707938 |
|  | Dental cement, Paladur powder | Kulzer | 64707963 |
|  | Dental cement, Super Bond 2000 | Sun Medical |  |
|  | Diazepam Injectable solution 10mg/2ml | Roche | Cip : 3400931112433 |
|  | DiIC18(3) | Interchim | 46804A |
|  | Neuro-Trace 500/5225 Green Fluorescent Nissl Stain, | Invitrogen | N21480 |
|  | Ketamine 1000 (1000mg/kg) | Virbac | GTIN : 03597132111010 |
|  | Lidocaine, 20mg/ml injectable solution | Aguettant | Cip : 3400936272606 |
|  | Meloxicam, Metacam 2mg/ml | Boehringer Ingelheim | GTIN : 04028691523598 |
|  | Pentobarbital, Euthasol | Dechra Veterinary Products | GTIN 08718469445110 |
|  | Pilocarpine hydrochloride | Sigma Aldrich | P6503 |
|  | Scopolamine methyl nitrate | Sigma Aldrich | S2250 |
|  | Sevoflurane Sevotek | Axience | GTIN 03760087152968 |
|  | Silicone gel | Dow Corning | 3-4680 |
|  | Sulfamethazine and trimethoprim, Amphoprim injectable solution | Virbac Animal Health | GTIN 03597132201094 |
|  | Tergazyme | Alconox |  |
|  | Xylazine, Rompun 2% | Bayer | GTIN : 04007221032311 |
| Surgery tools | Stereotaxic frame | Kopf | Model 963 |
|  | Motorized micromanipulator | Scientifica | IVM single |
|  | Volvere Vmax Dental Drill | NSK |  |
|  | Heating pad | Harvard apparatus | 55-7020 |
|  | MouseOx Pulse oximeter | Starr life sciences |  |
|  | Suture thread 4-0 Prolene | Ethicon | 4-0 Prolene |
|  | Cryostat | Microm microtech | NX50 |
| Implant elements and materials | 00-90 x 1/2 screw-oval fillister-brass | J.I.morris miniature fasteners | P0090B500 |
|  | 00-90 hex nut (brass) | J.I.morris miniature fasteners | N0090B |
|  | Form2 3D printer | Formlabs |  |
|  | Adhesive foam tape (20mm x 5m x 4mm) | RS pro | 554-866 |
|  | Copper tape | 3M | T118112 |
|  | Epoxy adhesive | RS components | 850-934 |
|  | Grey pro resin | Formlabs | RS-F2-PRGR-01 |
|  | Screws 00-90 3/32" (stainless steel) | Antrin Miniature Specialties Inc | B002SG89RA |
|  | Tulip pin ic socket | ASSMANN WSW | AR 06 HZL-TT |
|  | Form Cure UV chamber | Formlabs | FH-CU-01 |
|  | Wires (ground and reference) | Phoenix Wire Inc | 36744MHW |
| Acquisition rig | Cable | Imec |  |
|  | Cage | Tecniplast | 2000P00SU |
|  | Headstage | Imec |  |
|  | Neuropixels 1.0 probe | Imec |  |
|  | PXIe acquisition module | Imec |  |
| Software, Algorithm | Kilosort3 |  | https://github.com/MouseLand/Kilosort |
|  | Matlab R2021b (9.11) | Mathworks | https://www.mathworks.com/ |
|  | Open Ephys GUI | Open Ephys | https://open-ephys.org/gui |
|  | Phy |  | https://github.com/cortex-lab/phy |
|  | Python |  | https://www.python.org/ |
|  | Tinkercad | Autodesk | https://www.tinkercad.com |

